# Mobile antimicrobial resistance genes in probiotics

**DOI:** 10.1101/2021.05.04.442546

**Authors:** Adrienn Gréta Tóth, István Csabai, Maura Fiona Judge, Gergely Maróti, Ágnes Becsei, Sándor Spisák, Norbert Solymosi

## Abstract

Even though people around the world tend to consume probiotic products for their beneficial health effects on a daily basis, recently, concerns were outlined regarding the uptake and potential intestinal colonisation of the bacteria that they transfer. These bacteria are capable of executing horizontal gene transfer (HGT) which facilitates the movement of various genes, including antimicrobial resistance genes (ARGs), among the donor and recipient bacterial populations. Within our study, 47 shotgun sequencing datasets deriving from various probiotic samples (isolated strains and metagenomes) were bioinformatically analysed. We detected more than 70 ARGs, out of which *rpoB* mutants conferring resistance to rifampicin, *tet(W/N/W)* and potentially extended-spectrum beta-lactamase (ESBL) coding *TEM-116* were the most common. Numerous ARGs were associated with integrated mobile genetic elements, plasmids or phages promoting the HGT. Our findings raise clinical and public health concerns as the consumption of probiotic products may lead to the transfer of ARGs to human gut bacteria.

## Introduction

Probiotics and probiotic products have gained a worldwide reputation and popularity in our everyday lives irrespective of cultural background, geographic location or social standards. Beneficial health effects assigned to probiotics have been reported in several studies.^1^ What these studies have in common, is that they state that microbes carried in probiotics must remain present in the intestinal tract for a shorter or longer period of time to exert the expected beneficial effects. Nevertheless, the success of colonisation depends on several factors, thus the certainty of its realisation varies from individual to individual.^2^ Recently, however, the possibility of some unfavourable or sometimes even adverse effects of probiotic consumption have also been raised.^3^ Several publications indicate that bacterial strains included in probiotic compounds, powders and capsules may contain antimicrobial resistance genes (ARGs).^4–7^ Genes, including ARGs of the probiotic bacteria, can be transmitted to bacteria within the intestinal tract of the consumers by horizontal gene transfer (HGT). If such ARGs are received by pathogenic bacteria, the efficacy of antibiotic therapy prescribed as medical intervention for the diseases they cause may lessen. HGT can take place by transformation, conjugation or transduction. All these processes have one important property in common, namely, a DNA fragment getting introduced to a recipient cell. Apart from transformation, by which any gene can be taken up by the bacterium from its environment, the routes of HGT require special active delivery processes. By conjugation, cell-to-cell contact provides the opportunity for a copy of a plasmid to translocate to a recipient bacterium.^8^ In contrast, transduction negates the necessity for cell-to-cell contact, as in this case bacteriophages act as a conduit for shuttling genes among bacteria.^9^ The genetic environment of the genes, possibly ARGs, involved in the transfer, has a significant influence on the efficacy of the latter two HGT processes, i.e. on the genes’ mobility. The transferability of ARGs is facilitated by the presence of mobility genes in their tight genetic environment. If the genes harbour on plasmids or prophages, the chance of their transfer increases. By probiotics with supposedly mobile ARGs, the likelihood of gene transmission to other bacteria in the intestinal tract increases. Our study focuses on the ARG content and mobility in the metagenome of various probiotics and bacterial strains which were isolated from probiotics and analyzed using a unified bioinformatic approach.

## Results

The analysis of the sequencing data from 20 isolates and 27 metagenomic (multi-microorganism) samples (Table 1) is summarized in three sections. Following the presentation of the bacteriome and the identified AGRs (resistome), predictions regarding the mobility potential of ARGs were also summarized based on genetic characteristics that may play a significant role in HGT. If integrated mobile genetic elements (iMGE) are identified in the sequence context of an ARG, its greater mobility can be assumed. The case is the same if the contig harbouring an ARG are derived as plasmid or phage originated. In the mobilome section, we summarize these results.

**Table 1.**
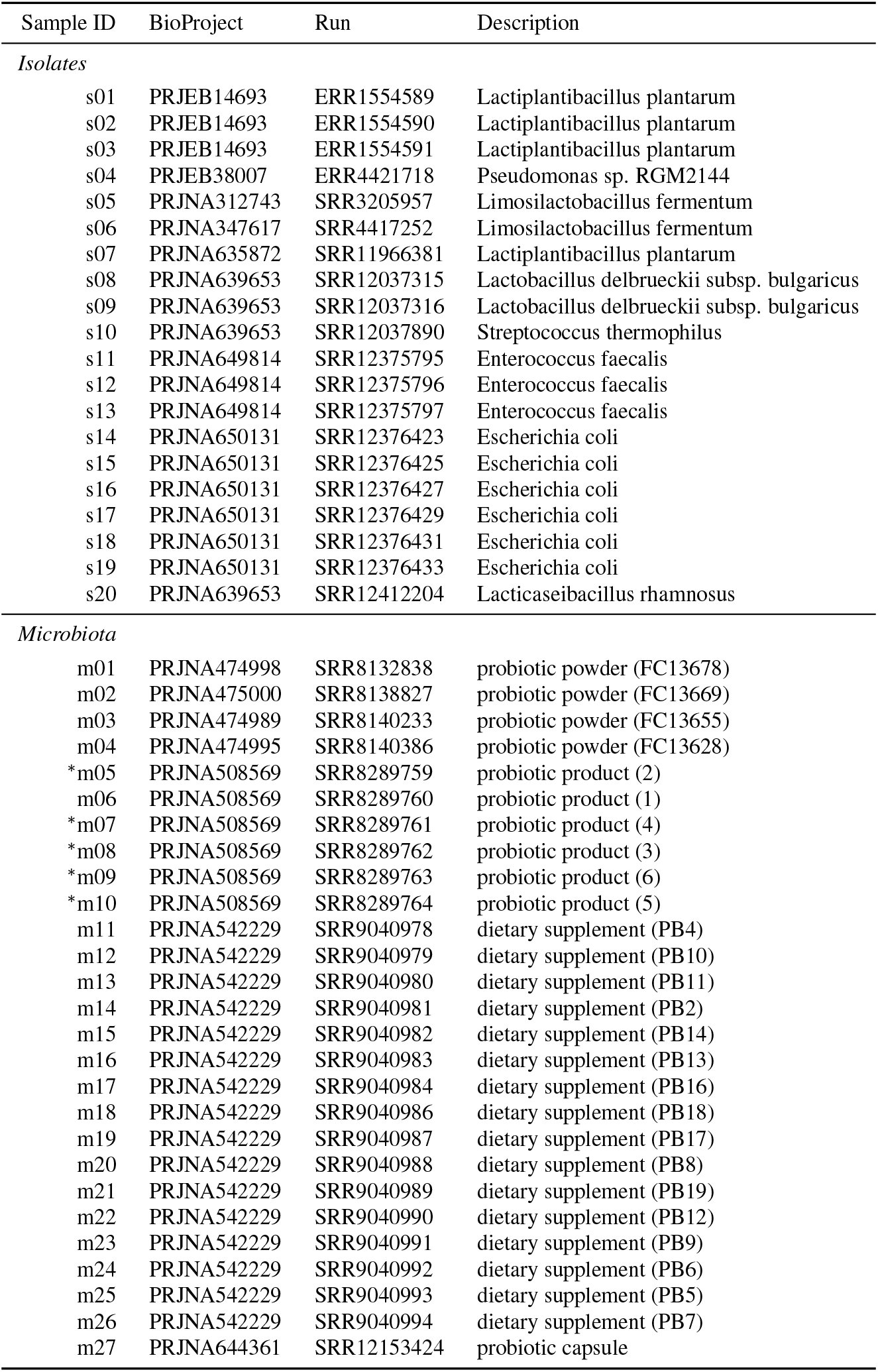
The list of analyzed samples obtained from NCBI SRA. In the unified names of the samples the first character corresponds to the type of the sample (s and m, isolate and metagenome, respectively), the second tag is a sequence number. Except the signed (^∗^) all samples were paired end sequenced. The last column shows the available information about the biosamples.

### Bacteriome

By taxon classification, the number of reads aligning to bacterial genomes differed in the various samples. The median bacterial read count of the metagenomic samples was 8.2 × 10^6^ (IQR: 4.4 × 10^6^). The median sequencing depth of the isolated strains was 220 (IQR: 94.8). The taxonomic origin of the short reads generated from isolates is shown in Table 1. The relative abundances of genera that achieved more than 1% of the bacterial hits in any of the metagenomic samples is shown in Figure 1. These dominant genera (with mean prevalence) in descending order were *Lactobacillus* (40%), *Enterococcus* (35%), *Bifidobacterium* (34%), *Limosilactobacillus* (34%), *Lactococcus* (32%), *Lacticaseibacillus* (31%), *Bacillus* (26%), *Weizmannia* (22%), Ligilactobacillus (19%), Streptococcus (18%), Lactiplantibacillus (12%), Sphingobacterium (2%).

**Figure 1.**
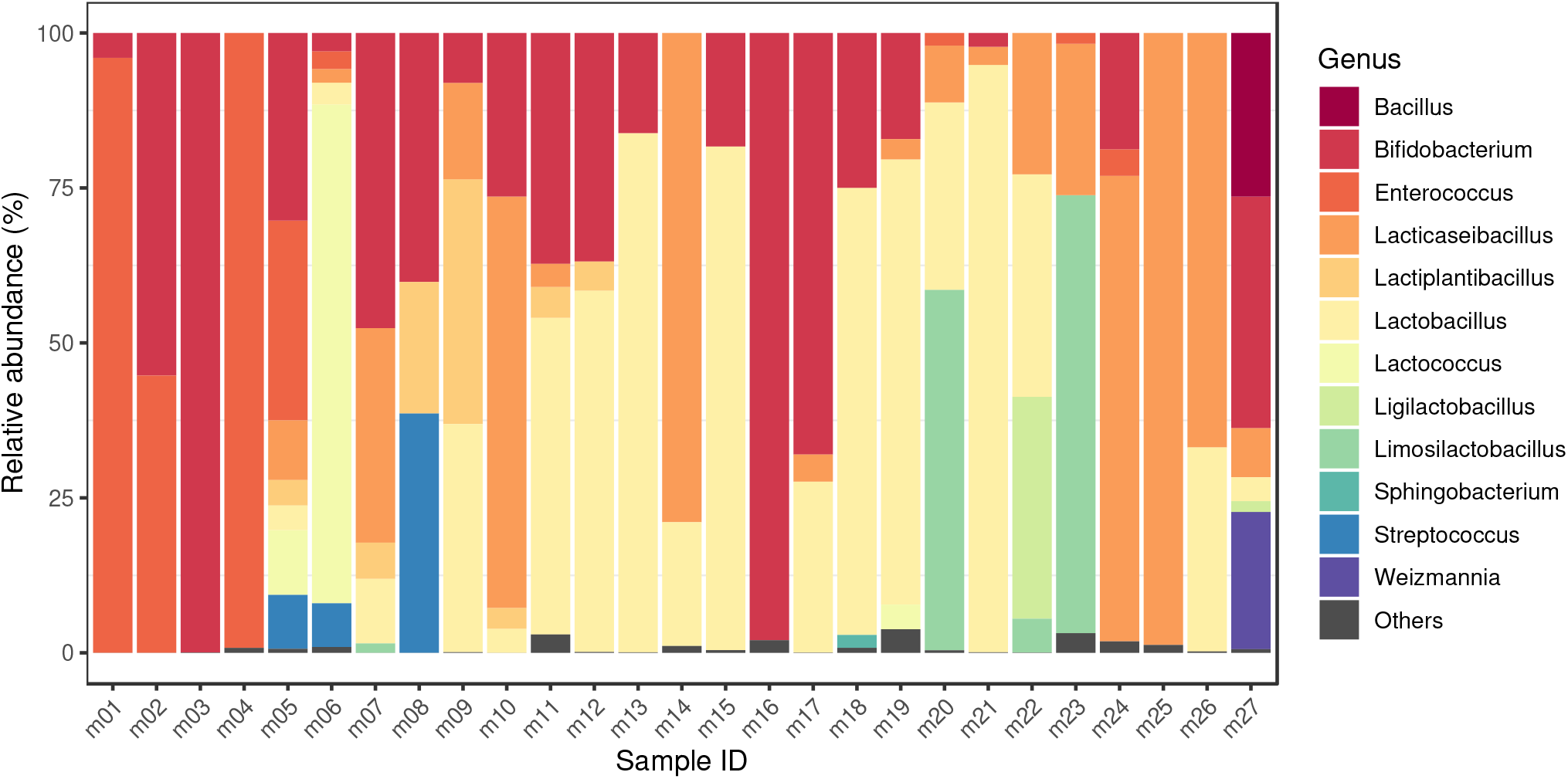
Bacteriome. The relative abundances of genera that achieved more than 1% of the bacterial hits in any of the metagenomic samples. The dominant genera (with mean prevalence) in descending order were *Lactobacillus* (40%), *Enterococcus* (35%), *Bifidobacterium* (34%), *Limosilactobacillus* (34%), *Lactococcus* (32%), *Lacticaseibacillus* (31%), Bacillus (26%), Weizmannia (22%), Ligilactobacillus (19%), Streptococcus (18%), Lactiplantibacillus (12%), *Sphingobacterium* (2%). Sample accession numbers for the Sample IDs are listed in Table 1.

### Resistome

The median length of the filtered contigs harbouring ARGs constructed by de novo assembly was 102,711 bp (IQR: 105,696). The number of ARGs found on the contigs ranged from 1 to 12. Besides 182 perfect ARG matches, further 225 hits were classified strict (RGI) and met the criteria of having 90% coverage and 90% sequential identity.

ARGs were detected in all metagenomic samples and in few isolates (Fig 2). The majority of isolates (s01, s02, s03, s04, s05, s06, s07, s08, s09, s10, s20) contained no ARG. The highest number of ARGs was found in samples s14-s19, obtained from sequencing six *Escherichia coli* strains isolated from the same probiotic product. It is important to highlight that we also found the *H-NS* gene in these samples which is not indicated in the figure, as its effect is anti-AMR. The most common ARGs were the *rpoB* mutants conferring resistance to rifampicin, *TEM-116* and *tet(W/N/W)* genes, detected in 18, 15 and 13 samples, respectively.

**Figure 2.**
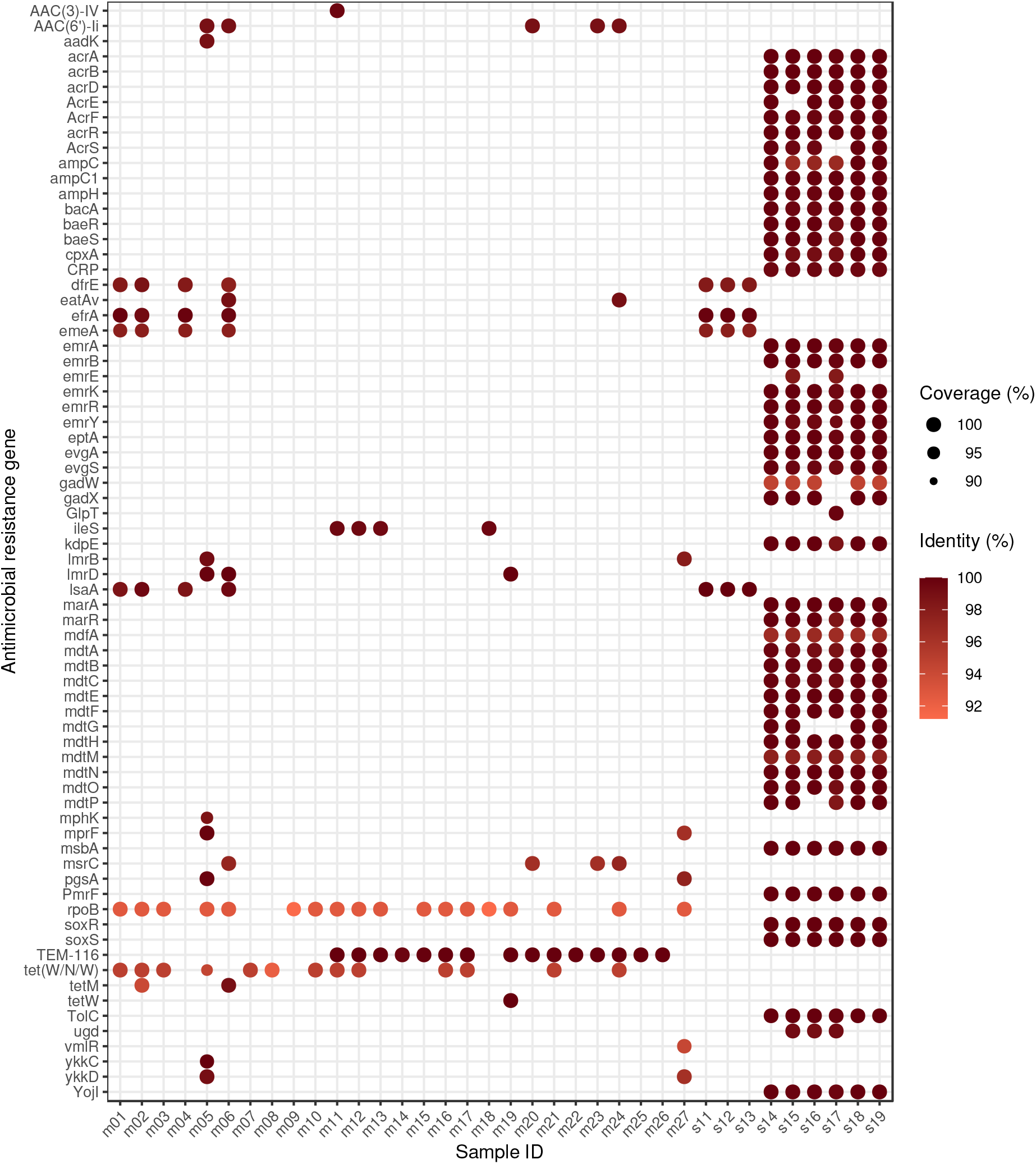
Identifed antimicrobial resistance genes (ARGs) by samples. For each sample-ARG combination, only the best finding is plotted. The size and the colour of the dots correspond to the coverage and the sequence identity of hits on reference genes, respectively. In sample s01-s10 and s20, there was no identifiable ARG. The gene names that are too long have been abbreviated (*acrA*: *Escherichia coli acrA*; *acrR*: *E. coli acrR* with mutation conferring multidrug antibiotic resistance; *ampC*: *E. coli ampC* beta-lactamase; *ampC1*: *E. coli ampC1* beta-lactamase; *ampH*: *E. coli ampH* beta-lactamase; *emrE*: *E. coli emrE*; *GlpT*: *E. coli GlpT* with mutation conferring resistance to fosfomycin; *ileS*: *Bifidobacterium bifidum ileS* conferring resistance to mupirocin; *marR*: *E. coli marR* mutant conferring antibiotic resistance; *mdfA*: *E. coli mdfA*; *mprF*: *Bacillus subtilis mprF*; *pgsA*: *B. subtilis pgsA* with mutation conferring resistance to daptomycin; *rpoB*: *Bifidobacterium adolescentis rpoB* mutants conferring resistance to rifampicin; *soxR*: *E. coli soxR* with mutation conferring antibiotic resistance; *soxS*: *E. coli soxS* with mutation conferring antibiotic resistance).

The proportion of resistance mechanisms was calculated based on the ARG diversity. The dominant mechanism of identified ARGs was the antibiotic efflux (58.33%), antibiotic inactivation (11.11%), antibiotic target alteration (11.11%), antibiotic target protection (9.72%), antibiotic target alteration and antibiotic efflux (4.17%), antibiotic efflux and reduced permeability to antibiotic (1.39%), antibiotic target alteration and antibiotic efflux and reduced permeability to antibiotic (1.39%), antibiotic target alteration and antibiotic target replacement (1.39%), antibiotic target replacement (1.39%).

There was no detectable ARG in the studied samples originating from Lacticaseibacillus rhamnosus, Lactiplantibacillus plantarum, Lactobacillus delbrueckii subsp. bulgaricus, Limosilactobacillus fermentum, Pseudomonas sp. RGM2144 or Streptococcus thermophilus species.

The identified ARGs associated with bacteria by species are as follows. *Bacillus subtilis*: *aadK*, *B. subtilis mprF*, *B. subtilis pgsA* with mutation conferring resistance to daptomycin, *bmr*, *lmrB*, *mphK*, *vmlR*, *ykkC*, *ykkD*. *Bifidobacterium animalis*: *B. adolescentis rpoB* mutants conferring resistance to rifampicin, *tet(W/N/W)*. *B. bifidum*: *B. adolescentis rpoB* mutants conferring resistance to rifampicin, *B. bifidum ileS* conferring resistance to mupirocin, *tet(W/N/W)*. *B. breve*: *B. adolescentis rpoB* mutants conferring resistance to rifampicin, *tetW*. *B. longum*: *B. adolescentis rpoB* mutants conferring resistance to rifampicin, *tet(W/N/W)*. *Enterococcus faecalis*: *dfrE*, *efrA*, *efrB*, *emeA*, *lsaA*, *tetM*. *E. faecium*: *AAC(6’)-Ii*, *eatAv*, *msrC*. *Escherichia coli*: *acrB*, *acrD*, *AcrE*, *AcrF*, *AcrS*, *bacA*, *baeR*, *baeS*, *cpxA*, *CRP*, *emrA*, *emrB*, *emrK*, *emrR*, *emrY*, *eptA*, *E. coli acrA*, *E. coli acrR* with mutation conferring multidrug antibiotic resistance, *E. coli ampC* beta-lactamase, *E. coli ampC1* beta-lactamase, *E. coli ampH* beta-lactamase, *E. coli emrE*, *E. coli GlpT* with mutation conferring resistance to fosfomycin, *E. coli marR* mutant conferring antibiotic resistance, *E. coli mdfA*, *E. coli soxR* with mutation conferring antibiotic resistance, *E. coli soxS* with mutation conferring antibiotic resistance, *evgA*, *evgS*, *gadW*, *gadX*, *kdpE*, *marA*, *mdtA*, *mdtB*, *mdtC*, *mdtE*, *mdtF*, *mdtG*, *mdtH*, *mdtM*, *mdtN*, *mdtO*, *mdtP*, *msbA*, *PmrF*, *TEM-116*, *TolC*, *ugd*, *YojI*. *Lactococcus lactis*: *lmrD*. *Streptomyces albulus*: *AAC(3)-IV*.

The ARGs belonging to the genome of *Bacillus subtilis* may play a role in the appearance of resistance against aminoglycosides, lincosamides, macrolides, oxazolidinones, peptides, phenicols, pleuromutilins, streptogramins, tetracyclines; *Bifidobacterium animalis*: rifamycins, tetracyclines; *Bifidobacterium bifidum*: mupirocins, rifamycins, tetracyclines; *Bifidobacterium breve*: rifamycins, tetracyclines; *Bifidobacterium longum*: rifamycins, tetracyclines; *Enterococcus faecalis*: acridine dye, diaminopyrimidines, fluoroquinolones, lincosamides, macrolides, oxazolidinones, phenicols, pleuromutilins, rifamycins, streptogramins, tetracyclines; *Enterococcus faecium*: aminoglycosides, lincosamides, macrolides, oxazolidinones, phenicols, pleuromutilins, streptogramins, tetracyclines; *Escherichia coli*: acridine dye, aminocoumarins, aminoglycosides, benzalkonium chlorides, carbapenems, cephalosporins, cephamycins, fluoroquinolones, fosfomycins, glycylcyclines, lincosamides, macrolides, monobactams, nitroimidazoles, nucleosides, penams, penems, peptides, phenicols, rhodamines, rifamycins, tetracyclines, triclosans; *Lactococcus lactis*: lincosamides; *Streptomyces albulus*: aminoglycosides.

### Mobilome

The frequencies of iMGEs, phages and plasmids associated with ARGs by bacteria of origin are summarized in Figure 3.

**Figure 3.**
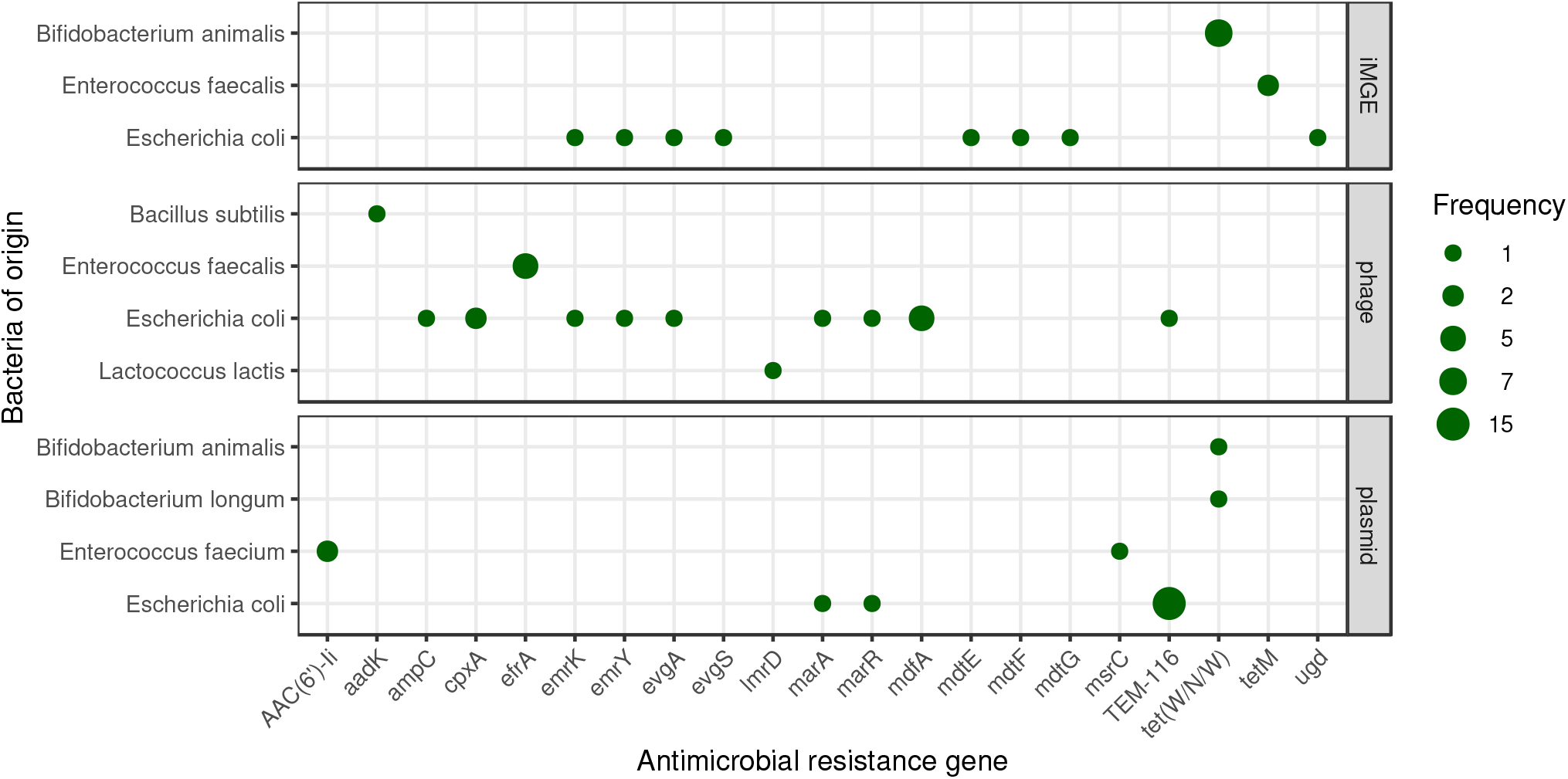
Mobile antimicrobial resistance gene frequency by bacteria of origin. The size of the dots indicates the occurrence frequency of the given gene flanked by iMGE, positioned in plasmid or phage.

### Coexistence of ARGs and iMGEs

Based on the distance method proposed by Johansson et al. (2021)^10^ iMGE associated ARGs were detected in three species (*Bifidobacterium animalis*, *Enterococcus faecalis* and *Escherichia coli*). In seven metagenomic samples (m01, m02, m03, m07, m16, m17, m24) we found tet(W/N/W) associated with ISBian1 insertion sequence on contigs classified as *B. animalis* originated. In two further samples (m02, m06) on *E. fecalis* originated contigs, *tetM* is linked to the transposon Tn6009. The ARG *mdtG* in the *E. coli* sample s14 and the ARG ugd in s15 are associated with IS3 and IS100 respectively. On two different contigs in the sample s17, multiple ARGs were detected with iMGE. One of them has the ISKpn24 associated with *mdtE* and *mdtF*. The other one has the IS102 linked to *emrY*, *emrK*, *evgA* and *evgS* genes. According to the average nucleotide identity (ANI) analysis most of the contig region of iMGE and associated ARGs had a high level of conservation (ANI>97%). Nevertheless, both contigs classified as *E. faecalis* originated showed ANIs below 80%.

### Plasmids

In the sample m08 and m21, we identified one-one plasmid associated contig with *tet(W/N/W)* classified as *Bifidobacterium longum* and *Bifidobacterium animalis* originated respectively. In the samples m20 and m23 on *Enterococcus faecium* classified contigs of plasmids *AAC(6’)-Ii* was detected. Further *E. faecium* classified contigs of the sample m23 contained the gene *msrC*. In the samples m11, m12, m13, m14, m15, m16, m17, m19, m20, m21, m22, m23, m24, m25 and m26, *Escherichia coli* originated contigs from plasmids harboured the gene *TEM-116*. In the *E. coli* isolate sample s15, one contig of plasmid had the *marA* and *marR* genes.

### Phages

By phage prediction, only dsDNAphages were detected. One contig, classified as *Bacillus subtilis* from the m05 metagenomic sample, contained prophage harbouring gene *aadK*. One prophage in predicted *Enterococcus faecalis* originated contig was found in sample m04 having gene *efrA*. The same content was detected in sample m01 on contigs classified as *E. faecalis*. All three *E. faecalis* isolates (s11, s12, s13) contained contigs harbouring the gene *efrA* within a prophage. In sample m17 one *E. coli* classified contig had the gene *TEM-116*, while a *Lactococcus lactis* classified one carried the gene *lmrD* within a prophage. All the *E. coli* isolates contained contigs with prophages harbouring ARG. In the sample s17 and s19 the *mdfA* gene is presented within a prophage. The sample s15 contains contigs harbouring prophage with the gene *marA*, *marR*. The sample s16 harbours contigs with prophage having genes *emrK*, *emrY*, *evgA*. The gene *ampC* was found in sample s15, while the gene *cpxA* in samples s14 and s18 within prophages.

## Discussion

The results presented demonstrate that the bacteria of probiotics may not only carry significant amounts of ARGs, but in numerous cases, those genes may also be mobile, thereby contributing to their spread to other bacteria and having possible consequences on the antibiotic treatment efficacy.

Bacterial genera identified in the metagenomic samples also appear in many probiotic related articles of the current international literature. Various species of *Bacilli*, *Bifidobacteria*, *Enterococci*, *Lacticaseibacilli*, *Lactiplantibacilli*, *Lactobacilli*, *Lactococci*, *Ligilactobacilli*, *Limosilactobacilli* and *Streptococci* are the core members of commercial probiotic bacterial communities.^11–20^ Two identified bacterial genera (*Sphingobacterium*, *Weizmannia*) in the various samples are less frequent probiotic components. The possibility of exploiting *Sphingobacteria* in probiotic foods was previously mentioned based on the characterization of flour and batter samples of sorghum and pearl millet.^21^ Members of the genus were detected by the high-throughput sequence analyses of fermented beverages.^22^ Probiotic *Weizmannia* species (e.g. former *Bacillus coagulans*) have recently been reclassified,^23^ and have an unquestionable probiotic significance.^24^

While at least one ARG was found in each metagenomic sample, less than half of the isolates contained any of them. No ARG was detected in *Lacticaseibacillus rhamnosus*, *Lactiplantibacillus plantarum*, *Lactobacillus delbrueckii subsp. bulgaricus*, *Limosilactobacillus fermentum*, *Pseudomonas sp. RGM2144* or *Streptococcus thermophilus*. Contigs originating from *Bacillus subtilis*, *Bifidobacterium animalis*, *Bifidobacterium bifidum*, *Bifidobacterium breve*, *Bifidobacterium longum*, *Enterococcus faecalis*, *Enterococcus faecium*, *Escherichia coli*, *Lactococcus lactis* and *Streptomyces albulus* each contained at least one ARG.

The available literature was screened to evaluate our findings and gain reliable knowledge of the ARGs that could have been attached to bacteria at the species level. All ARGs found in *Bacillus subtilis* (*aadK*, *B. subtilis mprF*, *B. subtilis pgsA* with mutation conferring resistance to daptomycin, *bmr*, *lmrB*, *mphK*, *vmlR*, *ykkC*, *ykkD*) have previously been identified in *B. subtilis* and many of them were even reported to be specific for this species or the *Bacillus* genus.^25–31^ In the *Bifidobacterium* genus, ARGs were associated with four species (*B. animalis*, *B. bifidum*, *B. breve* and *B. longum*). None of the *B. animalis*, *B. bifidum*, *B. breve* and *B. longum* related *B. adolescentis rpoB* mutants conferring resistance to rifampicin and *tet(W/N/W)* are specific for the identified species but both genes have previously been described in them.^6, 32–35^ *B. bifidum ileS* conferring resistance to mupirocin reported in *B. bifidum* supposedly cannot be exclusively linked to this species of the genus, but it had been identified in it before.^36^ Out of the *Enterococcus faecalis* deriving genes, *dfrE* was first identified in *E. faecalis*,^37^ but according to a recent study it is not exclusive to this species anymore.^38^ The genes *efrA* and *efrB* have been described in *E. faecalis* and *E. faecium*.^39, 40^ Gene *emeA* has only been identified in *E. faecalis* so far.^39^ Apart from *E. faecalis*, *lsaA* has been attached to *Streptococcus agalactiae*, while *tetM* appears in a broad spectrum of bacterial species.^41–45^ All three ARGs (*AAC(6’)-Ii*, *eatAv*, *msrC*) associated with *E. faecium* have been previously published as appearing in this species, and the first two are even specific for it.^46–49^ All ARGs originating from *Escherichia coli* (*acrB*, *acrD*, *AcrE*, *AcrF*, *AcrS*, *bacA*, *baeR*, *baeS*, *cpxA*, *CRP*, *emrA*, *emrB*, *emrK*, *emrR*, *emrY*, *eptA*, *E. coli acrA*, *E. coli acrR* with mutation conferring multidrug antibiotic resistance, *E. coli ampC* beta-lactamase, *E. coli ampC1* beta-lactamase, *E. coli ampH* beta-lactamase, *E. coli emrE*, *E. coli GlpT* with mutation conferring resistance to fosfomycin, *E. coli marR* mutant conferring antibiotic resistance, *E. coli mdfA*, *E. coli soxR* with mutation conferring antibiotic resistance, *E. coli soxS* with mutation conferring antibiotic resistance, *evgA*, *evgS*, *gadW*, *gadX*, *kdpE*, *marA*, *mdtA*, *mdtB*, *mdtC*, *mdtE*, *mdtF*, *mdtG*, *mdtH*, *mdtM*, *mdtN*, *mdtO*, *mdtP*, *msbA*, *PmrF*, *TEM-116*, TolC, ugd, YojI) have previously been described in this species and many of them are even specific to it, according to the Comprehensive Antibiotic Resistance Database (CARD).^50, 51^ Gene *lmrD*, the only ARG deriving from *Lactococcus lactis* has been identified in this species along with some others.^52, 53^ Even though *AAC(3)-IV* has been identified in several studies,^54, 55^ according to our knowledge this is the first time it has been detected in *Streptomyces albulus*.

Gene *TEM-116*, which is often referred to as a clinically significant extended-spectrum beta-lactamase (ESBLs), was the most frequently identified finding in our study. ESBLs are most commonly defined as the members of a ubiquitous enzyme family that is capable of conferring resistance to penicillins, first-, second- and third generation cephalosporins and aztreonam, and of being impeded by beta-lactamase inhibitors such as clavulanic acid.^56^ The 400 *TEM* variants that have been identified so far, can be disclosed in two clusters with one deriving from *TEM-1* (the first *TEM* protein to be described) and one linked to *TEM-116* as a progenitor.^57^ In line with our findigs, gene *TEM-116* is reported to be present worldwide harboring in the conjugative plasmids of a wide range of Gram-negative bacteria. Despite its wide geographical dissemination, establishment on multiple plasmids and centrality in the *TEM* family network indicating it is a naturally occurring enzyme with microbiologically proven ESBL characteristics,^58, 59^ some concerns have arisen about its designation, after the gene was found in non-ESBL producing *Klebsiella pneumoniae* strains.^60^ Moreover, commercial Taq polymerases used in PCRs may be contaminated with *bla_T EM_*_−116_ DNA which could lead to the erroneous identification of the gene in samples that do not actually contain it.^61, 62^ In our study, each sample in which this gene was detected originated from the same bioproject (PRJNA542229). As the samples come from different dietary supplements, one may interpret that this finding is an artefact or contamination as a consequence of some sample preparation steps. Nevertheless, as more detailed information on sample preparation is not available, this issue cannot be resolved.

As seen above and as described in other publications^63^ there is still a great deal of variation in details which need to be clarified by the interpretation of ARGs. Nevertheless, the suspicion that the identified ARGs may undermine the efficacy of several antibiotic classes, including acridine dye, aminocoumarins, aminoglycosides, benzalkonium chloride, carbapenems, cephalosporins, cephamycins, diaminopyrimidines, fluoroquinolones, fosfomycins, glycylcyclines, lincosamides, macrolides, monobactams, mupirocins, nitroimidazoles, nucleosides, oxazolidinones, penams, penems, peptides, phenicols, pleuromutilins, rhodamines, rifamycins, streptogramins, tetracyclines and triclosans raises some clinical considerations. According to the latest CDC report on antimicrobial use in the U.S., amoxicillin (penam), azithromycin (aminoglycoside), amoxicillin and clavunalic acid (penam, increased activity), cephalexin (cephalosporin) and doxycycline (tetracycline) are the most commonly administered compounds.^64^ Moreover, based on the latest WHO report on global antimicrobial use, amoxicillin (penam), ciprofloxacin (fluoroquinolon), sulphametoxazole and trimethoprim are the most commonly prescribed oral drugs and ceftriaxone (cephalosporin), gentamicin (aminogylcoside) and benzylpenicillin (penam) are the most commonly used parenteral compounds in 4 surveyed countries of the African Region. In 6 countries of the Region of the Americas, amoxicillin (penam), cefalexin (cephalosporin) and doxycycline (tetracycline) are the antibiotics with the highest oral consumption rates and ceftriaxone (cephalosporin), oxacillin (penam) and gentamicin (aminogylcoside) are the ones with the highest parenteral use. In the European Region, reports were made of 46 countries. Among orally administered antibiotics, amoxicillin (penam), amoxicillin and beta-lactamase inhibitors (penam, increased activity) and doxycycline (tetracycline) are the top 3 compounds, while ceftriaxone (cephalosporin), gentamicin (amynoglycoside), and cefazoline (cephalosporin) are the most common parenteral ones. Amoxicillin (penam), azithromycin (macrolide) and amoxicillin and beta-lactamase inhibitors (penam, increased activity) are the most commonly consumed oral antibiotics and ceftriaxone (cephalosporin), benzathine benzylpenicillin (penam) and procaine benzylpenicillin (penam) are the top 3 parenterally administered agents in the Eastern Mediterranean region. In the 6 surveyed countries of the Western Pacific Region amoxicillin (penam), doxycycline (tetracycline) and amoxicillin and beta-lactamase inhibitors (penam, increased activity) are the most commonly prescribed oral antibiotics, while cefazolin (cephalosporin), ceftriaxone (cephalosporin) and cefuroxime (cephalosporin) are the most frequently used parenteral compounds.^65^ Many of the most highly prioritized antibiotics could be affected by the presence of the detected ARGs. Meanwhile, out of the 15 antibiotic groups mentioned in the latest WHO report on critically important antimicrobials (CIA) for human medicine, 9 (aminoglycosides, carbapenems and other penems, cephalosporins, glycylcyclines, macrolides, monobactams, oxazolidinones, penicillins of various cathegories, quinolones) could possibly be affected by the ARGs identified in the various samples.^65^

It is important to underline that all the six *E. coli* isolates contained the gene *H-NS*, which plays a crucial role in the global gene regulation of various bacteria, including this species. The expression of a wide variety of genes is repressed by *H-NS*, and its deletion increases AMR and decreases drug accumulation. Even though this gene is stored in CARD,^50, 51^ its functional effect is adverse to that produced by ARGs.^66^

If ARGs are transmitted from probiotic bacteria to pathogenic bacteria within the consumer’s body, they may reduce the effectiveness of antibiotic therapy on the diseases participating pathogenic bacteria cause. The execution of gene transfer processes is more likely among bacteria that are in close physical proximity to each other and if the ARGs are associated to a mobile genetic environment. According to our results a considerable number of ARGs, such as those which are iMGEs-linked or have resided in plasmids or prophages.

The co-occurence of *tet(W/N/W)* and ISBian1 is in line with the findings of Rozman et al.,^6^ according to which all genomes of *B. animalis* (*subspecies lactis* or *animalis*) (n = 42) available in 2019 contained this gene. Moreover, by the investigation of the mobility characteristics of *tetW*, out of the transposases belonging to the family of the insertion sequences, ISBian1 seemed to be subspecies dependent in *B. animalis subsp. lactis* and flanking *tetW* in the majority of the strains.^6^ Our results of *tetM* linking to the transposon Tn6009 in *E. faecalis* is consistent with finding of Zangue et al. in South-African fecal samples.^67^

In two samples contigs harbouring *tet(W/N/W)* originating from *Bifidobacterium longum* and *Bifidobacterium animalis* were predicted to belong to plasmids. Several studies reported a wide prevalence of the *tetW* gene in *Bifidobacteria*.^6, 68–70^ While the co-occurrence of *tetW* and its flanking transposase is a common genetic feature of *B. animalis*, previous reports lack the identification of plasmids in *B. animalis*, even though the gene was associated with plasmids in other bacterial species.^71^ Despite *AAC(6’)-Ii* deriving from *E. faecium* being located in the chromosome in previous studies and it being defined as a chromosome-borne ARG on CARD^50, 51, 72^, our research indicates it took place in a plasmid according to our deliberations. An *E. faecium* associated contig contained gene *msrC*. According to the available literature, *msrC* is a chromosomal-encoded gene that is mentioned as an intrinsic property of *E. faecium* strains.^50, 51, 73^ While the expected bacterial species of origin was confirmed, our finding raises the likelihood of the gene being connected to a plasmid as well. In 15 samples, *E. coli* originated contigs harboured the gene *TEM-116*. Plasmid origin is a common feature of ESBL genes such as *TEM-116* according to several publications and is often referred to as a feature to facilitate their quick spread.^74–76^ In the *E. coli* isolate sample s15, one contig had the *marA* and *marR* genes. These widespread multiple antibiotic resistance genes had been identified on plasmids before.^77^ The gene *efrA* harbouring in contigs with a prediction of phage origins were identified in all publicly available *E. faecalis* genome sequences by Panthee and colleges too, along with a large set of phages in the genomes.^78^

An important aspect to take into consideration by the interpretation of the ARG occurrence in probiotics is that constituent strains can often naturally be, or rendered multiresistant, so that they can be co-administered with oral antibiotics and reduce gastrointestinal side effects.^79, 80^ In our study we could not distinguish whether the examined samples contained the ARGs for this purpose or not. Moreover, since ARGs were found in the vast majority of the samples tested, not a negligible proportion of them, it is possible that the presence of ARGs in bacteria may also play a role in their probiotic effect. ARGs play a role in defence against antibiotics and may provide general fitness against specific toxic effects for bacteria.^81, 82^ One may make an analogy with earlier practice. In livestock farming, antibiotics have been widely used as feed supplements for yield enhancement on a purely empirical basis. By this practice, antibiotics have put pressure on the gut bacteria and selected for resistant strains. As a result, animal feed efficiency and production indicators have improved. When probiotics are consumed, the expectation is that the “good” microorganisms, bacteria will colonize the gut. In numerous animal husbandry areas (e.g. broiler chicken production), the producers try to achieve this by continuous probiotic feeding. If these probiotics also contain bacterial strains harbouring ARGs, they achieve very similar results as before with the selective effect of antibiotic utilization. If it is true that certain ARGs are essential for the efficacy of probiotic bacteria, then the selection of strains should be carried out with consideration of the human health consequences. That is, bacterial strains that contain ARGs having no significant influence on human antimicrobial therapy efficiency should be used. However, based on our results, it can also be suggested that bacteria that do not contain ARGs at all can be used as probiotic components. To have a more detailed insight into this topic, several further studies would be needed. For instance they could also focus on reducing the mobility of genes whose presence may be necessary for the probiotic nature of particular bacteria. Based on the results, we consider it essential to monitor the ARG content of probiotic preparations and their mobility characteristics in the fight against antimicrobial resistance.

## Methods

### Data

The details of analyzed samples are listed in Table 1. One probiotic capsule was shotgun sequenced (PRJNA644361) by the authors. All further short read datasets were obtained from NCBI SRA repository.

### DNA extraction and metagenomics library preparation for PRJNA644361

Total metagenome DNA of the probiotic capsule sample was extracted using the UltraClean Microbial DNA Isolation kit from MoBio Laboratories. The quality of the isolated total metagenome DNA was checked using an Agilent Tapestation 2200 instrument. The DNA sample was used for in vitro fragment library preparation. In vitro fragment library way prepared using the NEBNext Ultra II DNA Library Prep Kit for Illumina. Paired-end fragment reads were generated on an Illumina NextSeq sequencer using TG NextSeq 500/550 High Output Kit v2 (300 cycles). Primary data analysis (base-calling) was carried out with Bbcl2fastq software (v2.17.1.14, Illumina).

### Bioinformatic analysis

Quality based filtering and trimming of the raw short reads was performed by TrimGalore (v.0.6.6, https://github.com/FelixKrueger/TrimGalore), setting 20 as a quality threshold. Only reads longer than 50 bp were retained and taxonomically classified using Kraken2 (v2.1.1)^83^ and a database created (24/03/2021) from the NCBI RefSeq complete archaeal, bacterial and viral genomes. For this taxon assignment the -`confidence 0.5` parameter was used to obtain more precise species level hits. The taxon classification data was managed in R^84^ using functions of the packages phyloseq^85^ and microbiome.^86^ The preprocessed reads were assembled to contigs by MEGAHIT (v1.2.9)^87^ using default settings. The contigs were also classified taxonomically by Kraken2 with the same database as above. From the contigs having more than 500 bp, all possible open reading frames (ORFs) were gathered by Prodigal (v2.6.3)^88^. The protein translated ORFs were aligned to the ARG sequences of the Comprehensive Antibiotic Resistance Database (CARD, v.3.1.1)^50, 51^ by Resistance Gene Identifier (RGI, v5.1.1) with Diamond^89^ The ORFs classified as perfect or strict were further filtered with 90% identity and 90% coverage. All nudged hits were excluded. The integrative mobile genetic element (iMGE) content of the ARG harbouring contigs was analyzed by MobileElementFinder (v1.0.3).^10^ Following the distance concept of Johansson et al.^10^ for each bacterial species, those with a distance threshold defined within iMGEs and ARGs were considered associated. In the MobileElementFinder database (v1.0.2) for *Escherichia coli*, the longest composite transposon (cTn) was the Tn1681. In the case of this species, its length (24,488 bp) was taken as the cut-off value. For *Lactococcus lactis*, this threshold was the length of the Tn5721 transposon, 11,256 bp. For *Enterococci*, the database contained cTn, the Tn6246 (5,147 bp) transposon, in *E. faecium* only. The same threshold was used for *E. faecalis* contigs. As the database neither contains species-level, nor genus-level cTn data for *Bacillus*, *Bifidobacterium* and *Streptomyces* species, a general cut-off value was chosen for the contigs of these species. This value was declared as the median of the longest cTns per species in the database (10,098 bp). The average nucleotide identity (ANI) was calculated for the region of iMGE and associated ARGs by FastANI (v1.32).^90^ The plasmid origin probability of the contigs was estimated by PlasFlow (v.1.1)^91^ The phage content of the assembled contigs was prediced by VirSorter2 (v2.2.1)^92^. The findings were filtered for dsDNAphages and ssDNAs. All data management procedures, analyses and plottings were performed in R environment (v4.0.4).^84^

## Acknowledgements

The authors would like to thank the providers of BioProjects (PRJEB14693, PRJEB38007, PRJNA312743, PRJNA347617, PRJNA474989, PRJNA474995, PRJNA474998, PRJNA475000, PRJNA508569, PRJNA542229, PRJNA577063, PRJNA635872, PRJNA639653, PRJNA649814, PRJNA650131). The research was supported by the TKP 2020-4.1.1.-TKP2020 of ITM Hungary, within the Digital Biomarker thematic programme of the Semmelweis University. It has also received funding from the European Union’s Horizon 2020 research and innovation program under Grant Agreement No. 874735 (VEO). GM received support from the Hungarian Academy of Sciences through the Lendület-Programme (LP2020-5/2020).

## Author contributions statement

NS takes responsibility for the integrity of the data and the accuracy of the data analysis. AGT, IC, NS and SS conceived the concept of the study. GM, NS and SS performed sample collection and procedures. AB, AGT and NS participated in the bioinformatic analysis. AB, AGT, GM, IC, MFJ and NS participated in the drafting of the manuscript. AB, AGT, GM, IC, MFJ, NS and SS carried out the critical revision of the manuscript for important intellectual content. All authors read and approved the final manuscript.

## Additional information

### Availability of data and material

The short read data of sample data are publicly available and can be accessed through the PRJEB14693, PRJEB38007, PRJNA312743, PRJNA347617, PRJNA474989, PRJNA474995, PRJNA474998, PRJNA475000, PRJNA508569,^93^ PRJNA542229, PRJNA577063, PRJNA635872, PRJNA639653, PRJNA644361, PRJNA649814, PRJNA650131 from the NCBI Sequence Read Archive (SRA).

### Ethics approval and consent to participate

Not applicable.

### Consent for publication

Not applicable.

